# Sexual dimorphism and allometry in human scapular morphology

**DOI:** 10.1101/2024.03.18.585525

**Authors:** Erin C.S. Lee, Rebekah L. Lawrence, Michael J. Rainbow

## Abstract

Scapular morphology is highly variable across the human population and appears to be sexually dimorphic – differing significantly between males and females. However, previous investigations of sexual dimorphism in scapula shape have not considered the effects of allometry (the relationship between size and shape). Disentangling allometry from sexual dimorphism is necessary because apparent sex-based differences in morphology could be due to inherent differences in body size. This study aimed to investigate sexual dimorphism in scapula shape and examine the role of allometry in sex-based variation. We used three-dimensional geometric morphometrics with Procrustes ANOVA to quantify scapula shape variation associated with sex and size in 125 scapulae. Scapular morphology significantly differed between males and females, and males tended to have larger scapulae than females for the same body height. We found that males and females exhibited distinct allometric relationships, and sexually dimorphic shape changes did not align with male- or female-specific allometry. A secondary test revealed that sexual dimorphism in scapula shape persisted between males and females of similar body heights. Overall, our findings indicate that sex-based differences in scapular shape are independent of size-shape relationships. Our results shed light on the potential role of sexual selection in human shoulder evolution, present new hypotheses for biomechanical differences in shoulder function between sexes, and identify relevant traits for improving sex classification accuracy in forensic analyses.

## Introduction

Scapular morphology is highly variable across the human population (Dwight, 1887; Graves, 1921). The multi-factorial sources of scapular shape variation are unclear; however, previous work has revealed significant differences in scapular shape between males and females (Maranho et al., 2022; Scholtz et al., 2010; Zdilla and Guzmán-López, 2023). Understanding sexual dimorphism in scapular morphology is important for contextualizing scapula features that are correlated with injury risk (Lee et al., 2020), developing personalized treatments, identifying relevant shape traits for classifying sex in forensic analyses (Atamtürk et al., 2019; Dabbs and Moore-Jansen, 2010; Di Vella et al., 1994; Er et al., 2020; Peckmann et al., 2016), and providing insights into the role of sexual selection in human evolution (Puts, 2010).

Previous studies identifying sexual dimorphism in scapular morphology have not corrected for the effects of allometry (the relationship between size and shape). Thus, it is unclear whether the observed differences are due to biological sex or size-shape relationships. Disentangling allometry from sexual dimorphism is necessary because males are, on average, larger than females. Therefore, apparent differences in morphology between sexes could be due to inherent differences in body size. This distinction is important. For example, from a human health perspective, if observed sex-based differences in scapular shape are used to inform sex-specific glenohumeral implants, but these differences are largely allometric (i.e.., due to size), then binning treatment into male and female options could disadvantage larger females and smaller males who fall outside of their group averages in size. In the knee, female-specific implants have not been shown to improve clinical outcomes (Johnson et al., 2011; Sappey-Marinier et al., 2020), and this could be because morphological sex differences do not persist after correcting for body height (Grelsamer et al., 2005) and femur length (Dargel et al., 2011).

It may be important to account for the scapula’s complex three-dimensional structure when examining sexual dimorphism and allometry. Previous work has found that 3D differences in morphology are associated with shoulder function across species (Lee et al., 2023; Young, 2006) and predictive of soft-tissue injury within humans (Lee et al., 2020). In forensic analyses of the scapula, sex classification models that incorporate multiplanar metrics yield better accuracy than models that consider only single-plane measurements (Dabbs and Moore-Jansen, 2010; Di Vella et al., 1994), indicating that 3D structure may differ between sexes. While these discriminative models use absolute (unscaled) metrics to achieve high classification accuracy, they do not parse out sexual dimorphism in shape from sexual dimorphism in size. Moreover, pre-determined discrete metrics may not capture 3D shape features important for function. Geometric morphometrics overcomes these limitations by analyzing landmarks across the entire bone, and scaling bones to a common size for shape comparison (Zelditch et al., 2012). Previous geometric morphometric studies of the scapula have quantified morphology using 2D projections. These studies have found that the male scapula tends to have more pronounced curvature on the medial and lateral borders, a more projected inferior angle, and a more concave glenoid compared to the female scapula (Maranho et al., 2022; Scholtz et al., 2010; Zdilla and Guzmán-López, 2023). However, the 2D approach could be missing important sexually dimorphic 3D features. Further, while all bones are scaled to a common size, these analyses did not account for the shape variation that persists due to size-shape relationships (allometry) even after scaling.

The purpose of this study was to investigate sexual dimorphism in scapular morphology and examine the role of allometry in sex-based shape variation. First, we tested for sexual dimorphism in three-dimensional scapular shape using landmark-based geometric morphometrics. We then determined the extents to which the observed sexual dimorphism is allometric and non-allometric. We hypothesized that males and females would exhibit different scapular morphology, and that observed differences would contain a significant non-allometric component.

## Methods

We combined three datasets (Kolz et al., 2021; Kolz CW, 2020; Lawrence et al., n.d.; Lee et al., 2020) for a total of 125 scapula meshes segmented from computed tomography scans. The datasets did not differentiate between sex and gender; therefore, we assumed that reported sex (male or female) was the sex assigned at birth (i.e. biological sex). There were 80 females and 45 males. The datasets included individuals varying in age (55 ± 10 years), condition of the rotator cuff (27.2% with full-thickness supraspinatus tears), and presence of shoulder symptoms (19.2% with symptoms). Since the sample’s observed shape variation could be associated with age and injury condition (Lee et al., 2020), we tested whether these traits differed between males and females in the sample. We performed two-proportions *Z*-tests for group differences in the proportion of full-thickness tears and presence of shoulder symptoms and a two-sample Welch’s *t*-test for differences in age.

Twenty-nine landmarks capturing articular site and muscle origin geometry were semi-automatically identified on each scapula in Checkpoint (*Stratovan Corporation*, Sacramento, CA) (Lee et al., 2020) (***Figure 1***). We performed Generalized Procrustes Analysis to scale and align the landmarks to a common size and orientation. This yielded 3D Procrustes coordinates that only describe variation in shape but retain allometric effects (shape variation explained by size).

**Figure 1:**
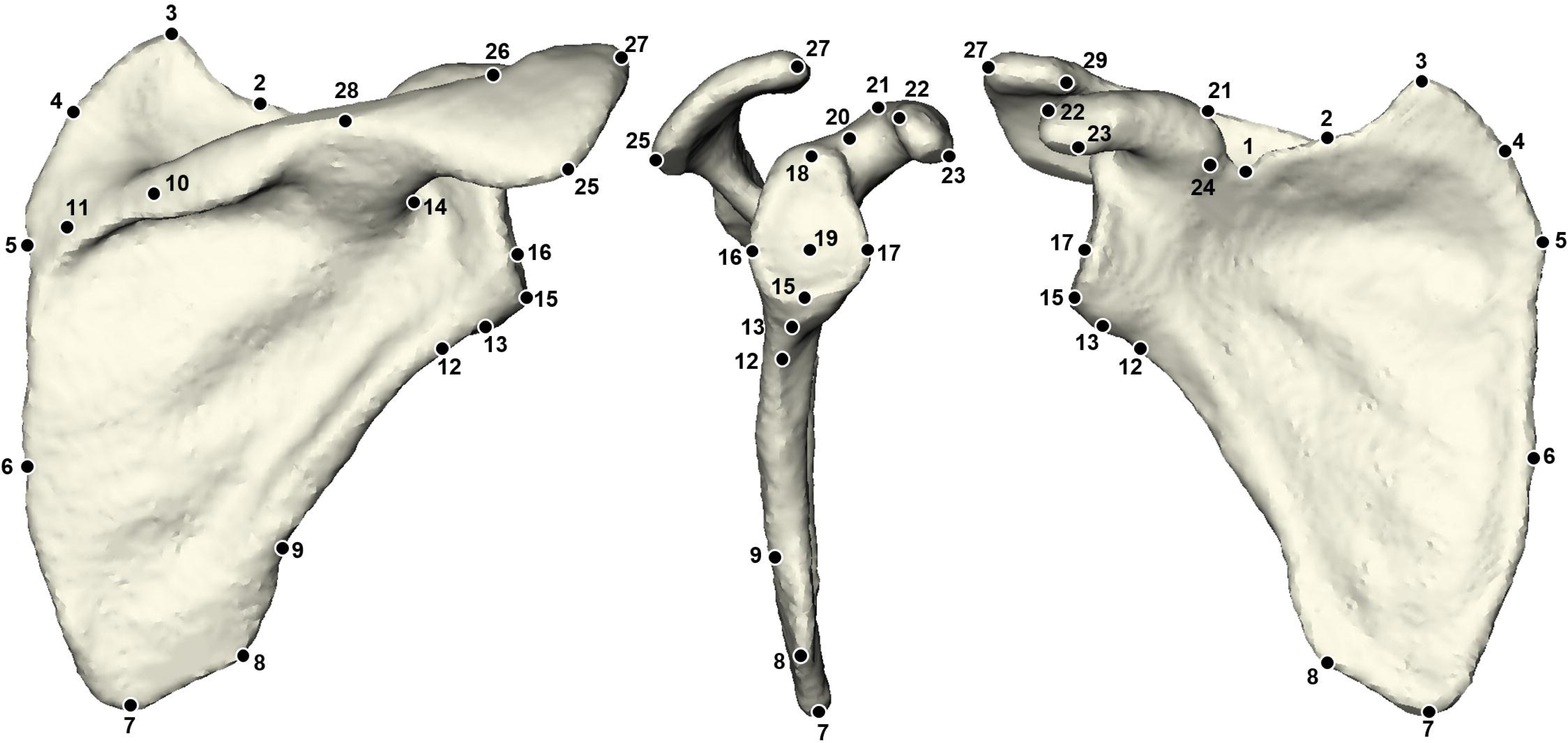
Scapula landmarks visualized on the individual with morphology closest to the sample mean. See Table 1 for landmark descriptions.

**Table 1:** Anatomical descriptions of each landmark. Adapted from (Lee et al., 2020)

| Landmarks |  |
| --- | --- |
| 1 | Suprascapular notch |
| 2 | Midpoint along superior border between (1) and (3) |
| 3 | Superior point on vertebral border |
| 4 | Midpoint between (3) and (5) along vertebral border |
| 5 | Intersection of spine with vertebral border |
| 6 | Midpoint between (5) and (7) on vertebral border |
| 7 | Inferior point on vertebral border |
| 8 | Superior lateral point of teres major fossa |
| 9 | Intersection of teres major fossa with axillary border |
| 10 | Deltoid tubercle (medial extent) |
| 11 | Confluence of scapular spine with body |
| 12 | Scapular neck |
| 13 | Triceps brachii insertion |
| 14 | Root of spine |
| 15 | Inferior extent of glenoid |
| 16 | Posterior extent of glenoid |
| 17 | Anterior extent of glenoid |
| 18 | Superior extent of glenoid |
| 19 | Shallowest point of glenoid along midline between (16) and (17) |
| 20 | Superior coracoid root on scapular body |
| 21 | Superomedial extent of proximal coracoid body |
| 22 | Superolateral extent of distal coracoid body |
| 23 | Inferolateral extent of distal coracoid body |
| 24 | Inferomedial extent of proximal coracoid body |
| 25 | Exterior angle of acromion |
| 26 | Interior angle of acromion |
| 27 | Lateral extent of distal acromion body |
| 28 | Transition between acromion and scapula spine |
| 29 | Medial extent of distal acromion body |

Body height was unknown for a subset of participants, so we quantified size using centroid size of the scapula (the square root of the sum of squared distances from each landmark to the centroid). To assess how scapula centroid size scaled with body size, we fit a linear regression model of centroid size on body height for participants with body height data (n = 115). We performed an Analysis of Covariance (ANCOVA) to examine whether the relationship between centroid size and body height differed between males and females.

We used Procrustes ANOVA to quantify the shape variation associated with sex and size (**Figure 2**). Briefly, Procrustes ANOVA begins with a multivariate regression of the 3D Procrustes coordinates (the *x*, *y*, and *z* locations of each landmark) on one predictor variable (Goodall, 1991; Zelditch et al., 2012). The multivariate regression model consists of a multi-component slope, where the components describe the change in each coordinate as a function of the predictor variable. We refer to this multi-component slope as the *shape vector*, which describes the overall change in 3D landmark configuration as a function of the predictor variable (Fischer and Mitteroecker, 2017). Unlike unsupervised dimension reduction techniques such as Principal Component Analysis (PCA), multivariate regression directly isolates shape features most relevant to the predictor variable of interest. Here, the predictor variables of interest were size and sex, yielding *allometry* and *sexual dimorphism* shape vectors, respectively.

**Figure 2:**
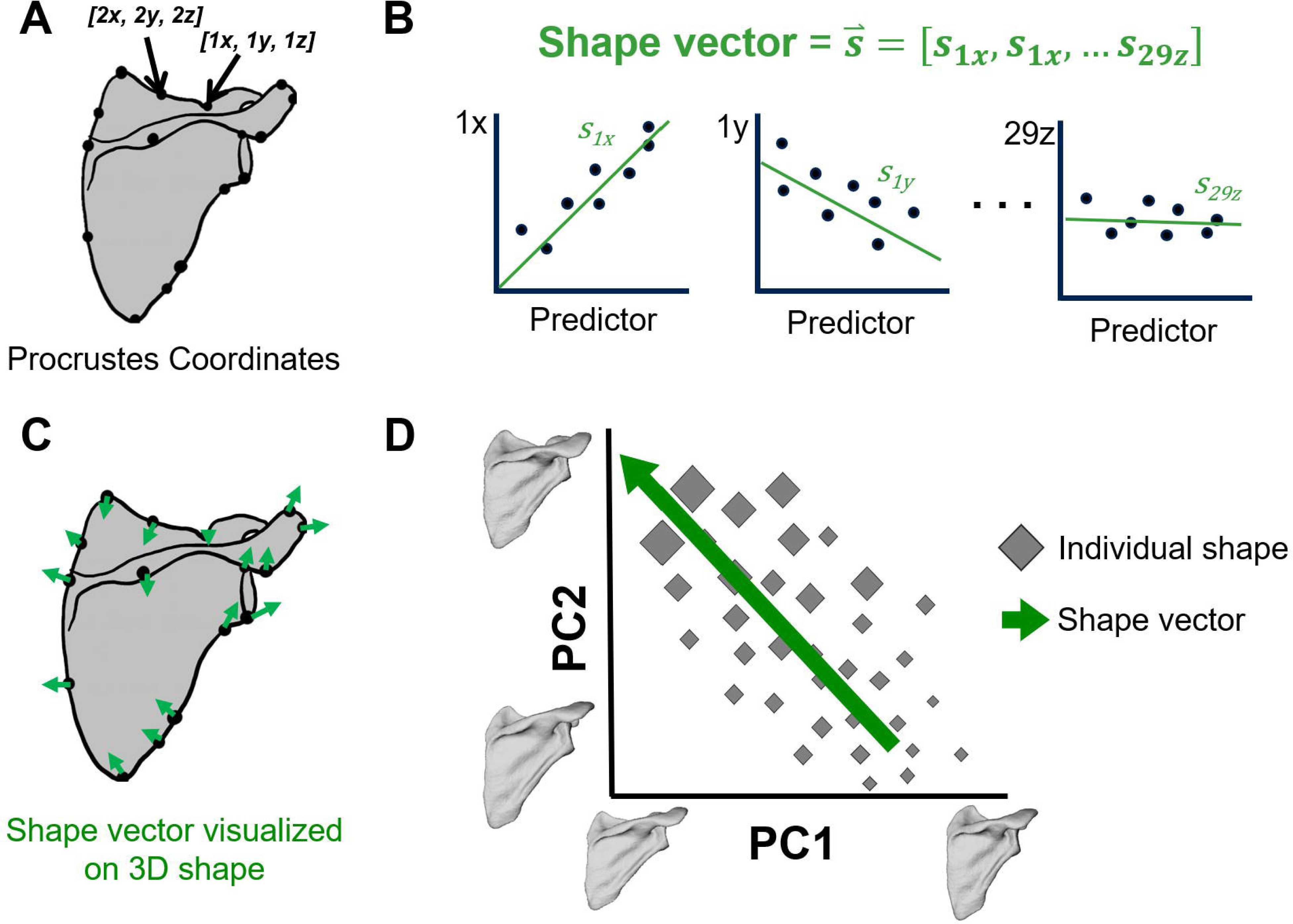
Visualization of a Procrustes ANOVA multivariate regression and the associated shape vector for a hypothetical relationship. (A) 3D Procrustes coordinates following scaling and alignment, visualized on one hypothetical shape. (B) The shape vector is equal to the slope of the multivariate regression model, where each component is the rate of change in the coordinate with respect to the predictor variable. Here, there are 29 three-dimensional landmarks, yielding 87 shape coordinates. (C) The shape vector visualized on the 3D shape, depicting the change in Procrustes landmarks for a unit increase in the predictor variable. (D) The simplified PC1-PC2 morphospace. The individual shapes comprising the hypothetical sample are scaled according to the magnitude of the predictor variable. The shape vector is oriented in the direction of increasing magnitude of the predictor variable.

We assessed the fit of the multivariate regression models using the distance-based approach characteristic of Procrustes ANOVA. In this approach, the residual variance is calculated as the Procrustes distance from the actual landmarks to the model-fitted landmarks of each scapula, effectively quantifying the total magnitude of shape variation not explained by the model (Goodall, 1991; Zelditch et al., 2012).

Randomized Residuals in a Permutation Procedure (RRPP) determined the statistical significance of each regression model, which we assessed at a 5% level of significance (Collyer and Adams, 2018). Procrustes ANOVAs were performed using the R packages *geomorph* (Adams et al., 2023) and *rrpp* (Collyer and Adams, 2023, 2018).

To visualize the shape vectors, we performed Principal Component Analysis (PCA) which identified the prevailing modes of scapular shape variation. We projected individual scapular shapes and shape vectors onto a simplified 2D shape space defined by Principal Component 1 (PC1) and Principal Component 2 (PC2) (**Figure 2**). While the PC1-PC2 projections aided in visualization, we did not analyze or compare the projected shapes or shape vectors since features affected by sexual dimorphism or allometry are not necessarily restricted to the first two PCs.

We first calculated the *sexual dimorphism vector* by regressing shape on sex using Procrustes ANOVA, where females were assigned a value of 0 and males were assigned a value of 1. The resulting sexual dimorphism vector was oriented from the mean female scapular shape to the mean male scapular shape. The sexual dimorphism vector alone does not provide any information on the extent to which the shape-sex relationship is driven by allometry because inherent differences in body size between sexes can contribute to apparent shape differences through allometric effects. Thus, we needed to quantify the shape variation associated with size. A *total allometry vector* – calculated by regressing shape on centroid size across the entire sample of males and females – could falsely capture an apparent shape-size relationship that is actually driven by sexual dimorphism in shape. Meaning, the shape differences observed between smaller and larger scapulae could be due to sex-based differences between female scapulae (which tend to be smaller) and male scapulae (which tend to be larger).

A common technique for disentangling allometry from sexual dimorphism is to correct the sexual dimorphism vector according to a pooled within-sex allometry regression (Klingenberg, 2016). This method captures the pooled relationship between size and shape across males and females while correcting for the inherent difference in body size between sexes. Importantly, the pooled within-sex allometry regression is only valid if the female- and male-specific allometric relationships are not significantly different (e.g., if size-related shape changes do not differ between sexes). To test this assumption, we computed male- and female-specific allometry vectors from separate Procrustes ANOVAs of shape coordinates on centroid size. We performed a pairwise comparison with 10000 RRPP permutations to test if male and female scapulae exhibited significantly different allometry vectors (i.e., different changes in shape as a function of centroid size) (Collyer and Adams, 2018). We calculated the angle between vectors to quantify the magnitude of the difference. Our test revealed that male and female allometry vectors differed significantly (θ = 86.1°, p = 0.02), violating the assumption required for a pooled within-sex allometry regression.

Therefore, we computed the angle between *the sexual dimorphism vector* and each *sex-specific allometry vector* as an alternate test to disentangle allometry from sexual dimorphism. If, for example, the female allometry vector is aligned with the sexual dimorphism vector (characterized by a small angle), it would indicate that as the female scapula gets larger, its shape gets closer to the mean male shape, suggesting that allometry is likely contributing to the observed sexual dimorphism.

We designed a vector comparison bootstrap test with 10000 iterations to assess the probability that the angle between the two test vectors under comparison (i.e., sexual dimorphism and within-sex allometry vector) exceeded the angle resulting from random variation (**Figure 3**). At each iteration, each test vector was recomputed from a multivariate regression on a bootstrap sample of the original data (with replacement). The iteration’s “test angle” was computed between the new bootstrap test vectors. We then calculated a “null angle” for each test vector, which described the angle between the bootstrap test vector and the original test vector. We computed the bootstrap test’s *p*-value as the proportion of iterations where one or both null angles exceeded the test angle. We implemented this test because angles computed between vectors with large dimensionality tend to be high. This vector comparison bootstrap test accounts for the vectors’ high-dimensionality and captures the goodness of fit of the Procrustes ANOVA regression models, since models with poorer fit would produce more variable bootstrap vectors.

**Figure 3:**
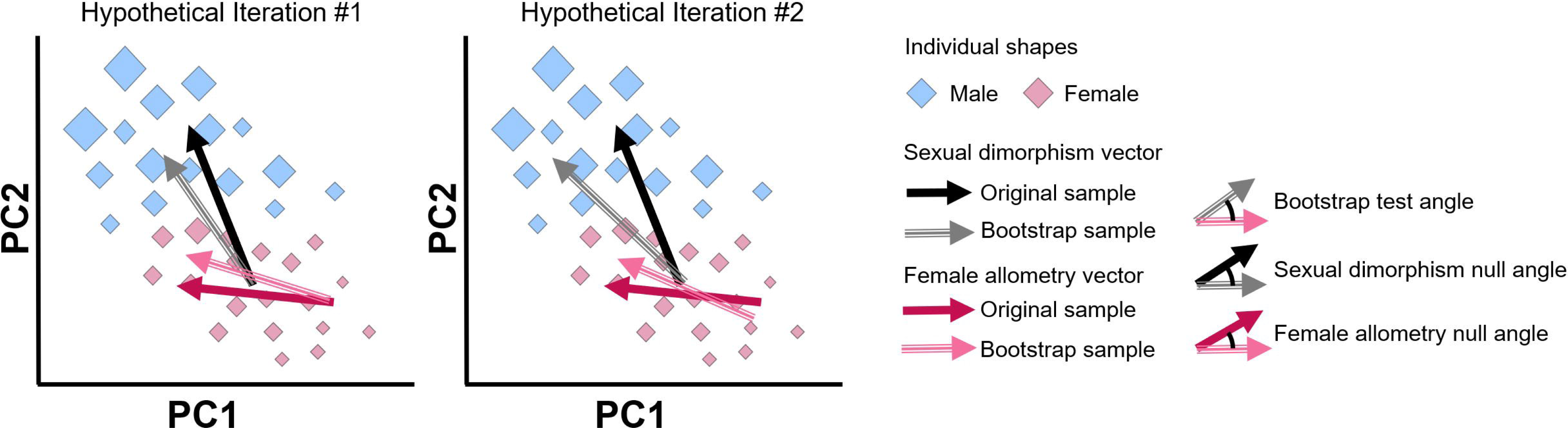
Vector comparison bootstrap test visualized for hypothetical data. The diamonds represent individual shapes and are scaled according to centroid size. The test angle was calculated between bootstrap samples of the two vectors being compared (here: sexual dimorphism and female allometry). Each null angle was calculated between the same vector recomputed with a bootstrap sample. Iteration #1 depicts an example where neither null angle exceeds the test angle. Iteration #2 depicts a case where the sexual dimorphism null angle exceeds the test angle.

We visualized the morphological changes described by each shape vector by warping the mesh closest to the mean (**Figure 1**) along the shape vector with a thin-plate spline. To visualize the shape changes associated with the sexual dimorphism vector, we generated warps of the mean male and female shapes. We also generated “exaggerated male” and “exaggerated female” shapes by extending the ends of the sexual dimorphism vector to three times its original length. The mean and exaggerated male and female 3D shapes are available as supplementary material. To visualize shape changes associated with the allometry vectors, we generated warps associated with the minimum and maximum centroid size observed in the sample.

We performed a secondary test to determine whether the scapular shape differed between males and females with the same body height. We identified an overlapping height range where the 5^th^ percentile of male heights defined the lower bound and the 95^th^ percentile of female heights defined the upper bound. We performed a Procrustes ANOVA of shape on sex for individuals with heights in the overlapping range.

## Results

The male and female groups did not differ significantly in age (p = 0.28, *t* = -1.1), the proportion with full-thickness tears (p = 0.76, *Χ*^2^ = 0.096), or the proportion experiencing shoulder symptoms (p = 0.56, *Χ*^2^ = 0.29).

Scapula centroid size was strongly correlated with body height (*R*^2^ = 0.72, p < 2.2e-16). ANCOVA revealed that the slope of centroid size on body height is not significantly different between males and females (p = 0.11), but the intercepts were significantly different (p < 1e-10). These results indicate that scapula size increases relative to body height at the same rate between males and females, but males tend to have larger scapulae than females for the same body height.

The sexual dimorphism vector significantly differentiated male and female scapulae (p < 0.001, *Z* = 4.66) but explained only a small amount of the total shape variation (*R*^2^ = 0.05) (**Figure 5**). The mean male scapula is superoinferiorly taller and mediolaterally narrower than the mean female scapula. The male glenoid is also larger, more retroverted, and less superiorly inclined than the female glenoid. The acromion and supraspinatus fossa are also antero-posteriorly broader in the male scapula than in the female scapula.

**Figure 4:**
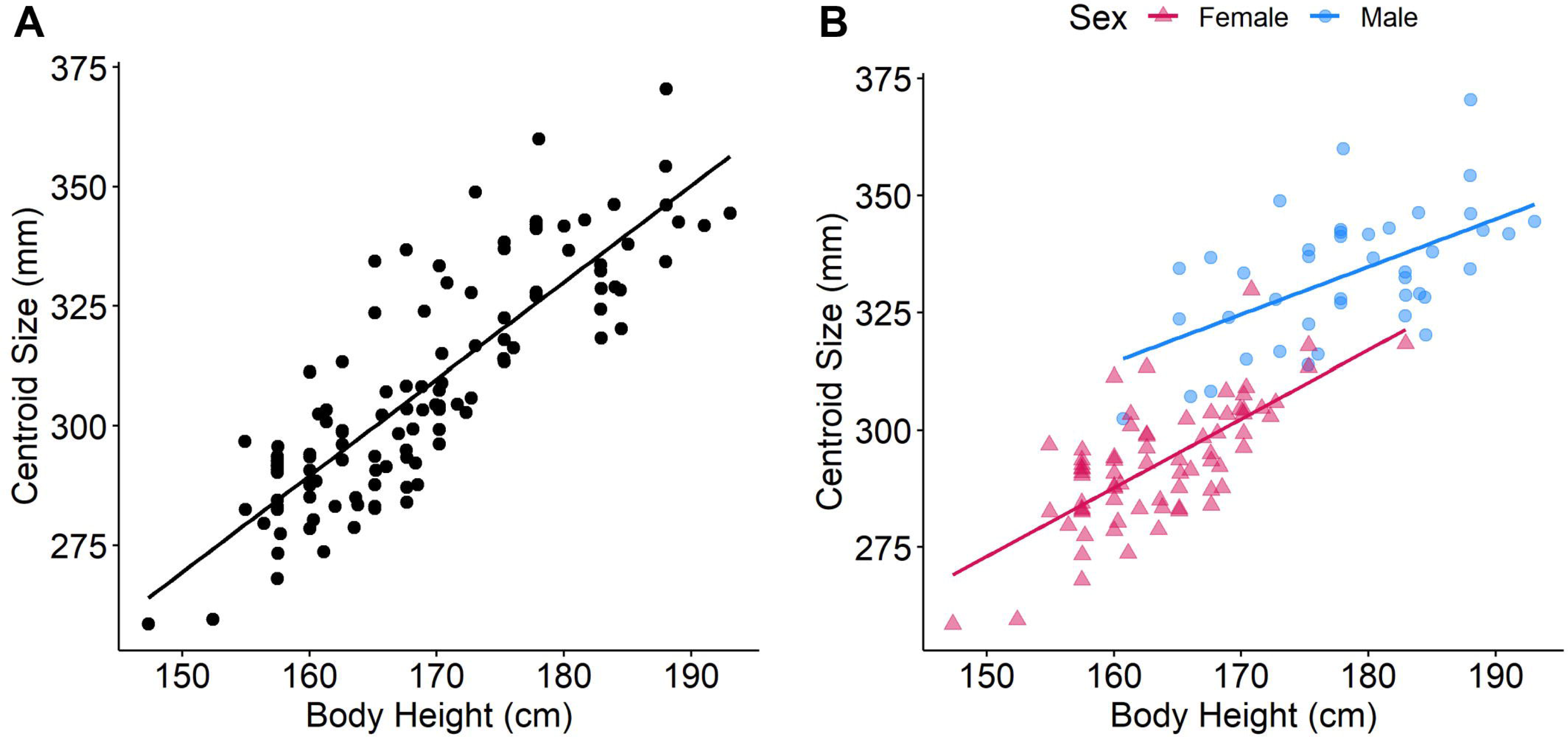
Scapula centroid size regressed against body height for individuals with body height information (n=115). (A) The black line indicates the pooled linear regression model. (B) The pink and blue lines indicate the sex-specific linear regression models.

**Figure 5:**
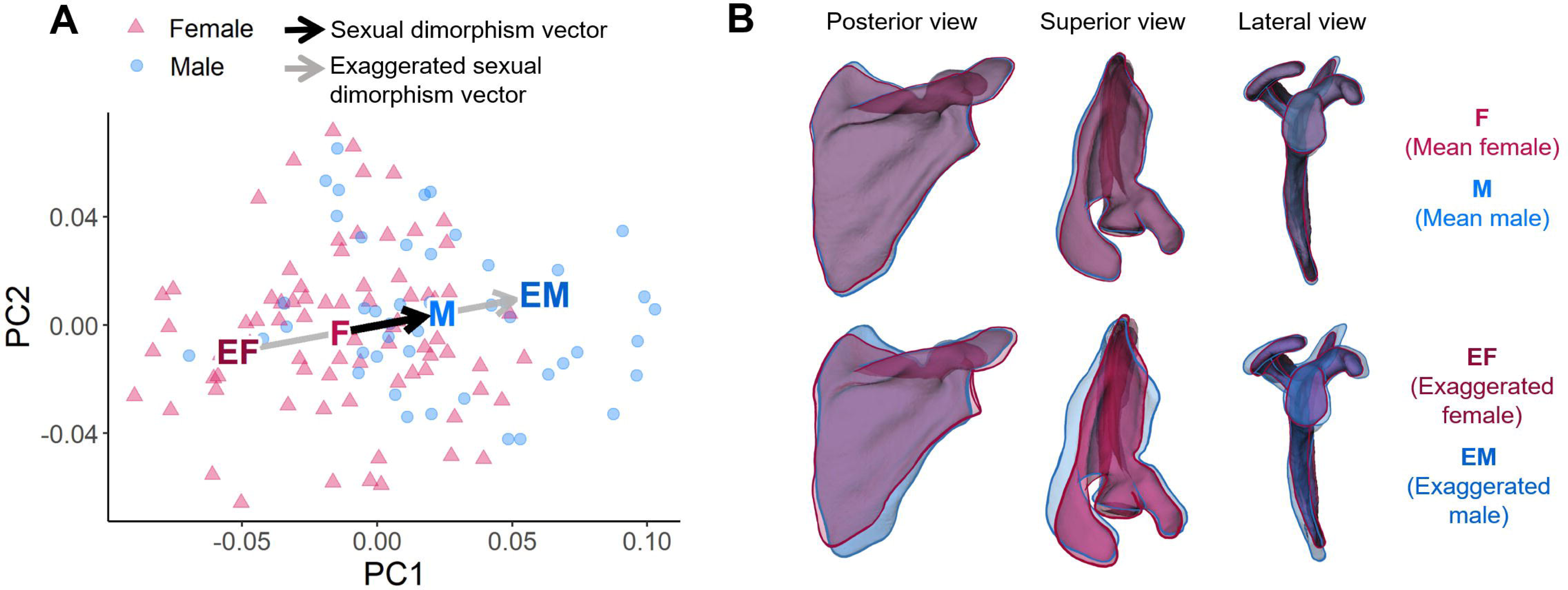
(A) The sexual dimorphism vector and individual shapes visualized on PC1-PC2 morphospace. The vector is oriented from the mean female shape to the mean male shape. (B) The difference in morphology visualized between the mean female and male shapes (top) and exaggerated female and male shapes (bottom).

Female and male allometric relationships were significantly different, as found in the previously violated assumption test for pooled regression (**Figure 6**). The female allometry vector was significant (p = 0.0013, *Z* = 2.90) but explained a small amount of total shape variation in the female sample (*R*^2^ = 0.031). Larger female scapulae tend to be superoinferiorly shorter and mediolaterally broader, exhibit smaller glenoids, and possess flatter acromions with reduced subacromial space. The male allometric relationship was not significant (p = 0.67, *Z* = -0.45, *R*^2^ = 0.018).

**Figure 6:**
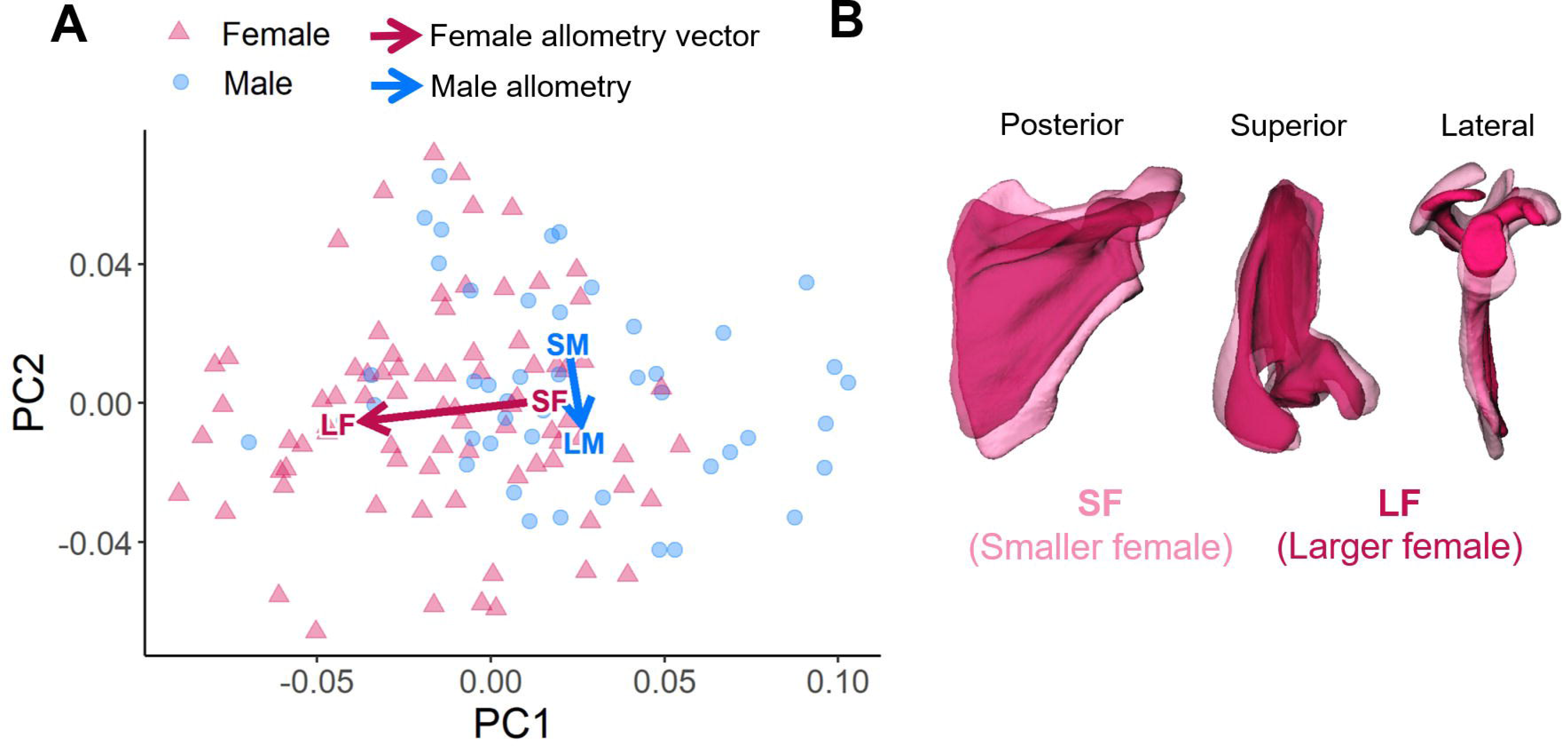
(A) The within-sex allometry vectors visualized on PC1-PC2 morphospace. The vectors begin at the shape associated with the minimum centroid size in the sample, and end at the shape associated with the maximum centroid size. (B) Female allometric relationship visualized on 3D morphology.

The vector comparison bootstrap test revealed that the sexual dimorphism vector differed significantly from the female allometry vector (mean ± SD angle from bootstrap tests: 120° ± 8°, p = 0) and the male allometry vector (89° ± 13°, p = 0.013, **Figure 7**).

**Figure 7:**
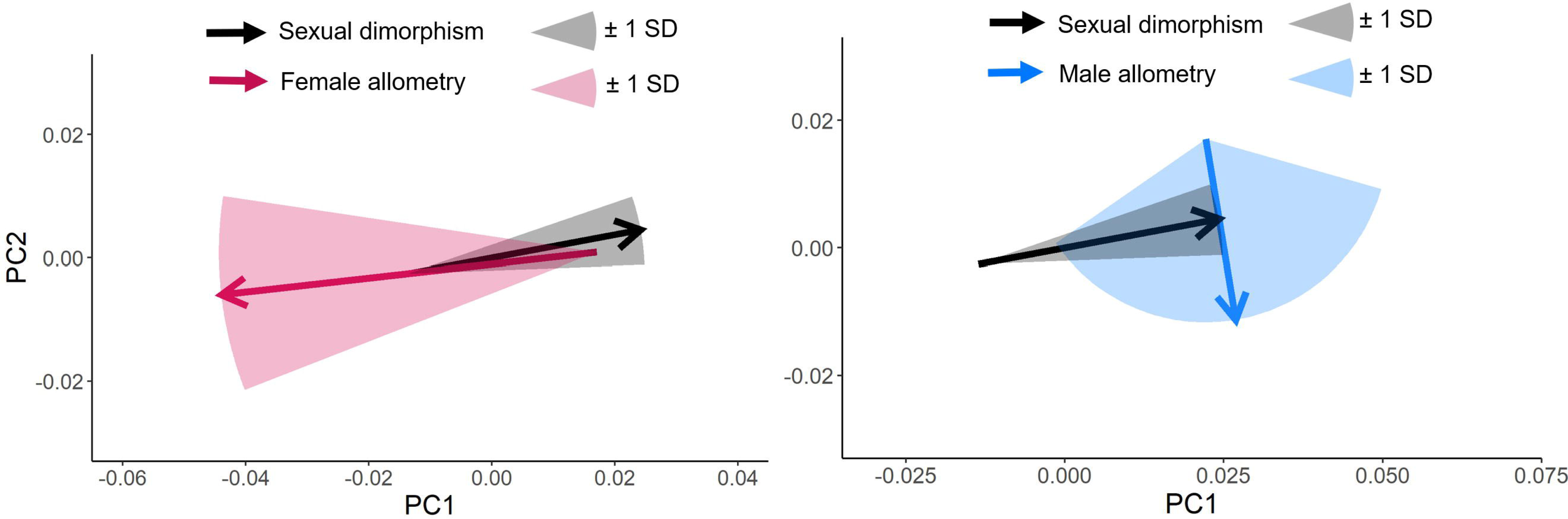
Variation in shape vectors from vector comparison bootstrap test. The shape vectors from the original sample are plotted on PC1-PC2 space, and shaded regions represent ± 1 standard deviation of the null angles (angle between bootstrap vector and original vector) projected onto PC1-PC2 space. (A) Comparison between sexual dimorphism and female allometry. (B) Comparison between sexual dimorphism and male allometry.

The overlapping height range (165.1 cm to 172.5 cm) included twenty-four females and six males (**Figure 8**). Sex differences persisted in this subsample, as the sexual dimorphism vector significantly differentiated female and male shapes (p = 0.0018, *Z* = 2.90, *R*^2^ = 0.091).

**Figure 8:**
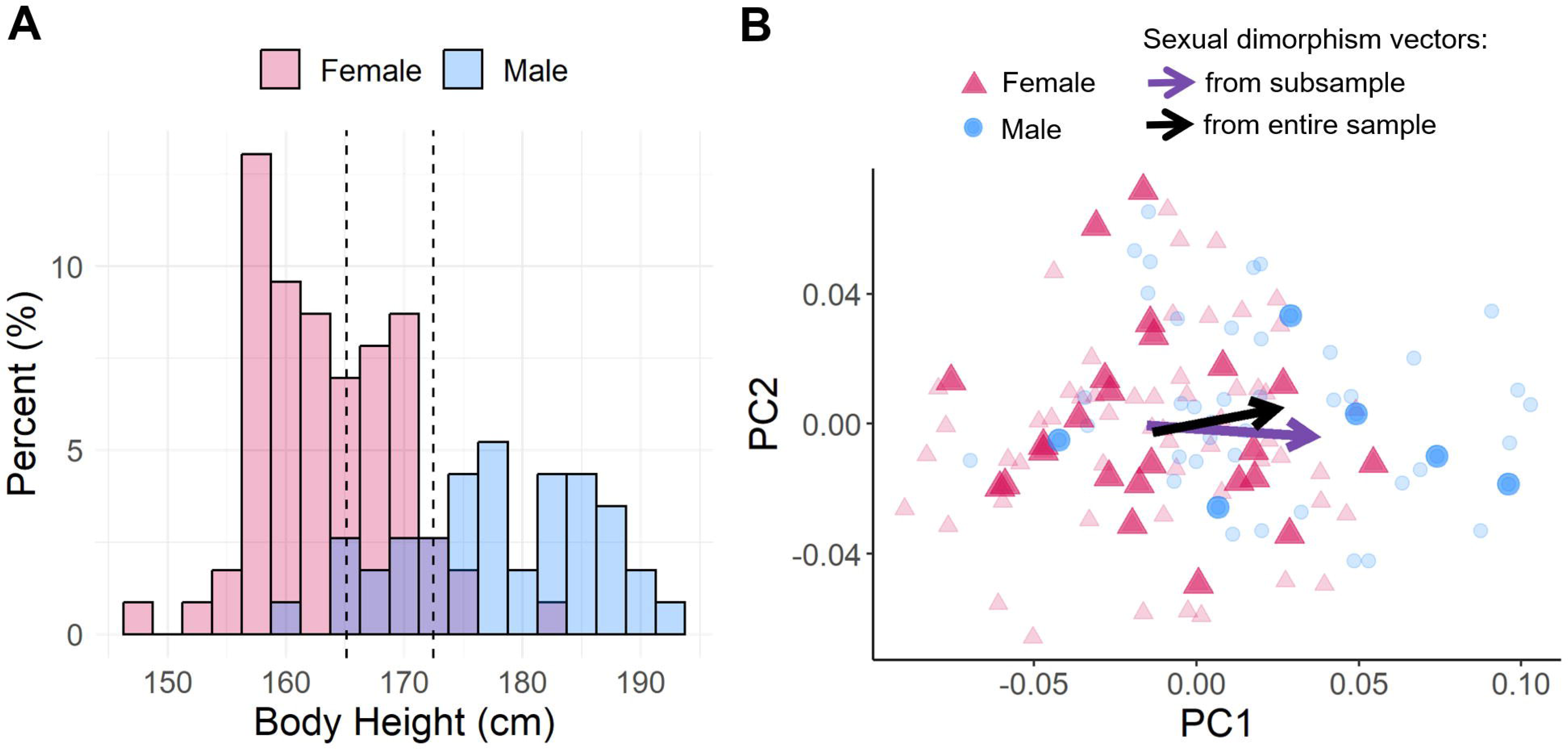
(A) Distribution of female and male body height. The dotted lines represent the lower bound (5^th^ percentile for males) and the upper bound (95^th^ percentile for females) that defined the range of overlapping height. (B) All individuals plotted on PC1-PC2 morphospace. The individuals included in the overlapping height range are represented by opaque and enlarged points while the remainder are transparent and small. The sexual dimorphism vector computed from the overlapping subsample is in purple.

## Discussion

In this study, we used three-dimensional geometric morphometrics to test for sexual dimorphism in scapular morphology and assessed the extent to which observed sexual dimorphism was explained by allometry (size-shape relationships). We identified scapula features that significantly differed between sexes. We also found that allometric relationships differed between males and females, and sexual dimorphic shape changes did not align with male- or female-specific allometry. Our findings indicate that sex-based differences in scapular shape are independent of size.

Although sex differences in scapular shape were significant, sex explained only 5% of the total shape variation (**Figure 5**). While this appears to be a small amount of shape variation, it is important to consider that small changes in shape variation may be associated with large functional differences. Our previous work identified that a principal component of scapular shape variation (PC4) explaining only 6.6% of the variation in overall shape was strongly associated with rotator cuff tears, which may imply that small changes in shape can lead to meaningful differences in shoulder function (Lee et al., 2020).

We were surprised to find that males and females exhibited distinct allometric relationships. We expected the size-related shape changes to reveal how features change in response to increasing loads associated with size. For example, limb bones scale allometrically across ungulate species - getting thicker relative to their length - to manage the stress associated with larger body mass (McMahon, 1975). The female allometric relationship was significant, with size explaining 3.1% of the total shape variation in females. Interestingly, the larger female scapulae had relatively smaller glenoids and smaller muscle attachment sites, which contradicts the changes expected to manage the higher joint contact stress and muscle mass expected with increased size.

Male-specific allometry, however, was non-significant and explained only 1.8% of the shape variation. This non-significant finding is consistent with previous work (Maranho et al., 2023) and could reflect the complex loading profile imposed on the scapula. Unlike in joints that consistently bear body weight, such as the knee, the forces applied to the scapula likely do not scale predictably with body size, and morphology may accommodate larger forces in various ways. However, given that allometry was significant in females, the lack of relationship could be due to the smaller male sample size. Future studies with a larger sample size should investigate the discrepancy in allometry observed between females and males. Nevertheless, our result suggests that investigations into allometry in human skeletal structures should account for potential differences in sex-specific relationships.

There are several potential sources of non-allometric sexual dimorphism in scapular morphology. For example, sexual dimorphism could reflect the role of sexual selection in human evolution (Puts, 2010). Researchers have proposed that sexual selection on fighting performance in male-male competition led to male-biased upper body specializations in apes (Morris et al., 2019a). Males have relatively higher muscle mass than women, with most of the added muscle allocated to the upper body (Lassek and Gaulin, 2009), and biomechanical testing suggests that males possess greater potential for powerful forward punching (Morris et al., 2019b). Sexual selection could explain male-associated shape characteristics such as the superoinferiorly-broader blade shape that increases the breadth of supraspinatus, subscapularis, and infraspinatus attachment sites. It could also explain why the male scapula is larger than the female scapula for the same body height (**Figure 4**). A larger scapula could accommodate muscle attachment sites for males’ proportionally larger upper body muscle mass.

Sexual dimorphism in scapular morphology could also arise from morphological integration between the scapula and pelvis (Young, 2004). The human pelvis exhibits substantial sexual dimorphism due to obstetric demands and allometric effects (Fischer and Mitteroecker, 2017). The shoulder and pelvic girdles demonstrate evolutionary covariance in quadrupedal primates (Agosto and Auerbach, 2021); thus, changes in shoulder morphology could be associated with changes in pelvic girdle morphology. However, the scapula and pelvis exhibit distinct evolutionary histories and genetic pathways (Young et al., 2019), so the potential for within-species sex-based covariance between the two structures is unclear.

Finally, the scapula may undergo plastic morphological changes in response to differences in muscle forces imposed by an individual’s activities and environment (Kuhns, 1945; Wolffson, 1950). Gender norms influence occupation and recreational activity (Akerlof and Kranton, 2000; Kane, 1990); thus, differences in activity – particularly during youth – could contribute to sexual dimorphism in shape. However, given we did not collect or analyze participants’ gender identity or physical histories, we cannot assume that plastic response to activity contributed to the sex-differences observed in our sample.

The sex differences in morphology identified here could have biomechanical implications for shoulder function and injury risk. Specific scapula features are associated with musculoskeletal diseases including rotator cuff tears and osteoarthritis (Banas et al., 1995; Bigliani et al., 1986; Hughes et al., 2003; Kim et al., 2012; Moor et al., 2013; Nyffeler et al., 2006; Pandey et al., 2016; Tétreault et al., 2004).

Biomechanical studies have shed light on how these “at-risk” shape features may alter shoulder joint mechanics in ways that are relevant for pathology, such as by altering muscle moment arms (Lee et al., 2020), rotator cuff tendon load (Gerber et al., 2014), and glenohumeral joint reaction force (Viehöfer et al., 2015). There is evidence that the prevalence and incidence of self-reported shoulder pain are higher in women than men (Bot et al., 2005; Greving et al., 2012; van der Windt et al., 1995), and women are at a higher risk of occupational repetitive strain injuries in the shoulder (Ashbury, 1995). While the causes are undoubtedly multifactorial, it is possible that sexual dimorphism in scapular shape alters biomechanical function in a manner that increases risk of shoulder pathology in females.

It is interesting to note that many characteristics of the female scapula are consistent with injury-associated shape differences including a narrower supraspinatus fossa (Lee et al., 2020), a laterally-projected acromion (Nyffeler et al., 2006), a superior-inclined glenoid (Hughes et al., 2003), and a combination of the latter two to yield a higher critical shoulder angle (Moor et al., 2013). For example, the female scapula is superoinferiorly shorter and mediolaterally broader than the male scapula. This blade shape results in narrower attachment sites for the supraspinatus, infraspinatus, and subscapularis and likely alters the muscles’ lines of action and potential for stabilizing the humerus head. The female glenoid is also superiorly inclined and anteverted. A superiorly inclined glenoid has been theorized to predispose the humeral head to superior migration and increase load on the supraspinatus (Hughes et al., 2003), and an anteverted glenoid is thought to increase stress on the posterior rotator cuff (Tétreault et al., 2004). Future computational and physical modelling should examine whether the observed sex differences contribute to rotator cuff pathomechanics.

Given that the variation explained by sex was small, any consideration of patient-specific implants should consider an individual’s unique anatomy, given that overall bone shape is not well predicted by sex alone. Future work should investigate alternate sources of three-dimensional shape variation to understand the factors influencing the scapula’s highly variable structure.

Our results could help inform discriminative models for sex classification in forensic analysis. In the absence of intact cranial and pelvic skeletal remains (Mays and Cox, 2000), the scapula is considered a feasible alternative for classifying an individual’s sex (Er et al., 2020). Consistent with previous studies, we found that the male scapula is disproportionately larger than the female scapula, confirming that models based on absolute (unscaled) linear metrics can take advantage of both sexual dimorphism in size and shape to improve classification accuracy. In addition, our shape analysis on scaled bones revealed metrics that are relatively larger on the male scapula. These included the vertebral border height, the distance between the posterior acromion and anterior coracoid, supraspinatus fossa breadth, and glenoid width. Therefore, including these unscaled metrics, as opposed to measures that are relatively larger in females – such as the medial-lateral breadth of the scapular blade and lateral acromion extension – could improve sex classification accuracy.

Our study had several limitations. First, our combined dataset was limited to participants from North America but scapular morphology and patterns of sexual dimorphism appear to be dependent on geographical location (Peckmann et al., 2016). Therefore, our results may not be generalizable to a global population. Second, our main analysis used scapula centroid size as a proxy for body size. Centroid size was strongly correlated with body height in a subsample of our dataset (**Figure 4**), but it may be more informative to investigate allometry with respect to body height directly to understand the relationship between shape and stature. While centroid size is less biased in quantifying size than single linear measures (such as vertebral border height), it is still affected by the landmark set and inherently dependent on shape (see the Pinocchio Effect (Klingenberg, 2021)). For these reasons, we performed a secondary analysis to test for sexual dimorphism in an overlapping height range. Although this range included a small sample of males and females, sexual dimorphism persisted in scapulae belonging to individuals of similar heights (**Figure 8**). Finally, our geometric morphometrics analysis used 29 discrete landmarks. The landmarks were semi-automatically identified and subject to user error. To mitigate inter-observer error, a single observer applied all landmarks across all participants for consistency. In addition, the sparse landmark set does not capture morphological complexities such as the curvature of the fossae or glenoid. Surface-based models or dense 3D semi-landmarks could overcome this limitation (Clouthier et al., 2023; Verhaegen et al., 2021). Currently, automatically establishing point-to-point landmark correspondences across the scapula is notoriously difficult due to its extreme curvature and thin scapular blade; however, rapidly developing methods may enable this approach in future analyses (Cates et al., 2017; Porto et al., 2021).

In this study, we used geometric morphometric to analyze sexual dimorphism in scapular shape while accounting for allometric effects. We observed sex-based differences that are not attributable to differences in size between males and females. Our findings shed light on the potential role of sexual selection in shaping male shoulder morphology, present new hypotheses for biomechanical differences in shoulder function between sexes, and offer insights for improving sex classification accuracy in forensic analyses.

## Supporting information

Mean and exaggerated male and female scapula meshes

## Acknowledgements

We thank Michael J. Bey (Henry Ford Health) and Heath B. Henninger (University of Utah) for contributing scapula meshes for the shape analysis. The authors have no conflicts of interest to declare.

## Author Contributions

Erin C.S. Lee contributed to the study concept and design, digitized landmarks, performed data analysis and interpretation, and drafted the original manuscript. Rebekah L. Lawrence contributed to data acquisition and provided critical revision of the manuscript. Michael J. Rainbow contributed to the study concept and design, assisted in drafting the original manuscript, and provided critical revision of the manuscript. All authors approved the final version of the manuscript.

## Supplementary Material

Three-dimensional bone models of the mean and exaggerated male and female scapulae are provided as STL files.

